# Gene expression analysis and proximity labeling reveal posttranscriptional functions of the yeast RNA Polymerase II regulator Def1

**DOI:** 10.1101/2024.09.16.613278

**Authors:** Oluwasegun T. Akinniyi, Shardul Kulkarni, Mikayla M. Hribal, Cheryl A. Keller, Belinda M. Giardine, Joseph C. Reese

## Abstract

Def1 is a yeast protein that promotes transcription elongation and regulates the degradation of RNA polymerase II during transcription stress. Although Def1 is localized in the cytoplasm, its functions in this cellular compartment are not yet understood. Despite its well-established roles in transcription, a comprehensive genome-wide analysis of its impact on gene expression has not been conducted. Here, we performed RNA sequencing (RNA-seq) analysis on cells lacking *DEF1* and surprisingly found that only a few hundred genes exhibited altered expression, both upregulated and downregulated. To evaluate mRNA synthesis and decay rates in these *DEF1*-deficient cells, we used a nascent transcription metabolic labeling technique called RATE-SEQ. As expected, we observed reduced synthesis rates across the genome in these cells. Additionally, a global decrease in mRNA decay rates was observed, suggesting that Def1 plays a role in the post-transcriptional regulation of mRNAs. The changes in synthesis and decay rates showed a strong correlation, indicating that this compensation helps buffer steady-state mRNA levels. To gain further insight into Def1’s functions, we conducted proximity labeling experiments to identify its protein binding partners within the cells. Our findings revealed that Def1 primarily interacts with cytoplasmic regulators involved in post-transcriptional processes, including proteins responsible for deadenylation, decapping, and translation regulation. Using an mRNA decay reporter assay, we demonstrated that recruiting Def1 to mRNA reduces its expression and accelerates its turnover. In summary, we have identified a novel cytoplasmic function for Def1, establishing it as a key regulator of gene expression in both transcription and mRNA decay.

## Introduction

Controlling gene expression involves coordinating multiple processes in both the cytoplasm and the nucleus. One important aspect of this coordination is known as gene expression buffering. This phenomenon occurs when changes in mRNA synthesis within the nucleus are offset by corresponding changes in mRNA decay in the cytoplasm (1–4). For example, complexes and proteins that were recognized for their impact on transcription in the nucleus can also influence post-transcriptional events in the cytoplasm (2, 3, 5). These proteins are compelling candidates for investigating how various stages of gene expression are co-regulated. How the coordination between processes is achieved is unclear, but it involves either the shuttling of these regulators between the nucleus and cytoplasm or indirectly through feedback loops.

DNA-damaging agents disrupt multiple steps in gene expression, affecting transcription, mRNA degradation, and protein synthesis (translation). One cellular response to damaged DNA is the ubiquitylation and subsequent degradation of the large subunit of RNA Polymerase II (RNAPII), known as Rpb1 (6–9). A key protein involved in this process is yeast <u>De</u>gradation <u>F</u>actor 1 (DEF1), which was first identified as a protein that co-purifies with the chromatin-bound transcription-coupled repair protein Rad26 (10). Def1 binds to arrested RNAPII and serves as a scaffold, facilitating the recruitment of ubiquitin ligases and other factors necessary for the degradation of RNAPII (6, 9, 11). Def1 undergoes proteasome-mediated proteolytic processing in the cytoplasm, which is dependent on DNA damage and transcription stress (12). This processing separates the N-terminus of the protein from its C-terminus, which is rich in polyglutamine and helps retain the protein in the cytoplasm. Def1 also plays roles in the nucleus, such as transcription initiation, elongation, telomere maintenance, and various DNA repair pathways (6, 13). Remarkably, despite its established role as a regulator of RNAPII, comprehensive gene expression analyses in *DEF1* mutants have not yet been reported.

While much attention has focused on Def1’s nuclear functions, it is an abundant protein found in the cytoplasm. A key question arises: does Def1 have functions in the cytoplasm, or is its presence in this compartment merely a means of preventing interference with RNAPII transcription in the nucleus under non-stress conditions? Evidence supporting the former hypothesis comes from proximity labeling (BioID) experiments, which have shown that subunits of the Ccr4-Not deadenylase complex interact with Def1 (14). Additionally, the presumed human homolog of Def1, UBPA2/2L, has been found to bind to multiple cytoplasmic RNA-binding proteins (15).

In this study, we performed gene expression analysis in conjunction with proximity labeling techniques to investigate the potential cytoplasmic functions of the yeast protein Def1. Our findings reveal that Def1 interacts with several cytoplasmic proteins involved in the post-transcriptional regulation of mRNAs, particularly those regulating mRNA decay. The absence of Def1 results in a significant decrease in mRNA decay rates globally. Importantly, when Def1 is recruited to an mRNA, it accelerates the turnover of that mRNA and suppresses its expression. Therefore, our results suggest that Def1’s presence in the cytoplasm is essential not only for preventing interference with transcription in the nucleus but also for regulating mRNA stability and expression.

## Results

### Def1 affects the steady-state RNA levels of a few transcripts

Def1 plays a vital role in regulating transcription elongation, particularly by removing stalled RNA polymerase II during transcriptional stress (6, 9, 12). Despite its regulatory function on RNAPII activity, no gene expression analyses of DEF1 mutants have been reported. To address this, we conducted RNA sequencing (RNA-seq) on wild-type and *def1Δ* mutant strains. Anticipating that Def1 would have widespread effects on transcription, we included *S. pombe* cells as a spike-in control before RNA isolation (see Experimental procedures). Two replicates of the experiment were analyzed, and displayed a high degree of correlation (R² > 0.99) as shown in Supplementary Fig. 1A. A total of 6,486 RNAs were detected, encompassing all types of RNA. After normalizing the data against the *S. pombe* spike-in reads, we identified differentially expressed genes (DEGs) using a cutoff of >2-fold change and a p_adj_ < 0.01. This analysis yielded 612 transcripts that were upregulated and 475 transcripts that were downregulated (Fig. 1A and Supplementary File 1).

**Figure 1:**
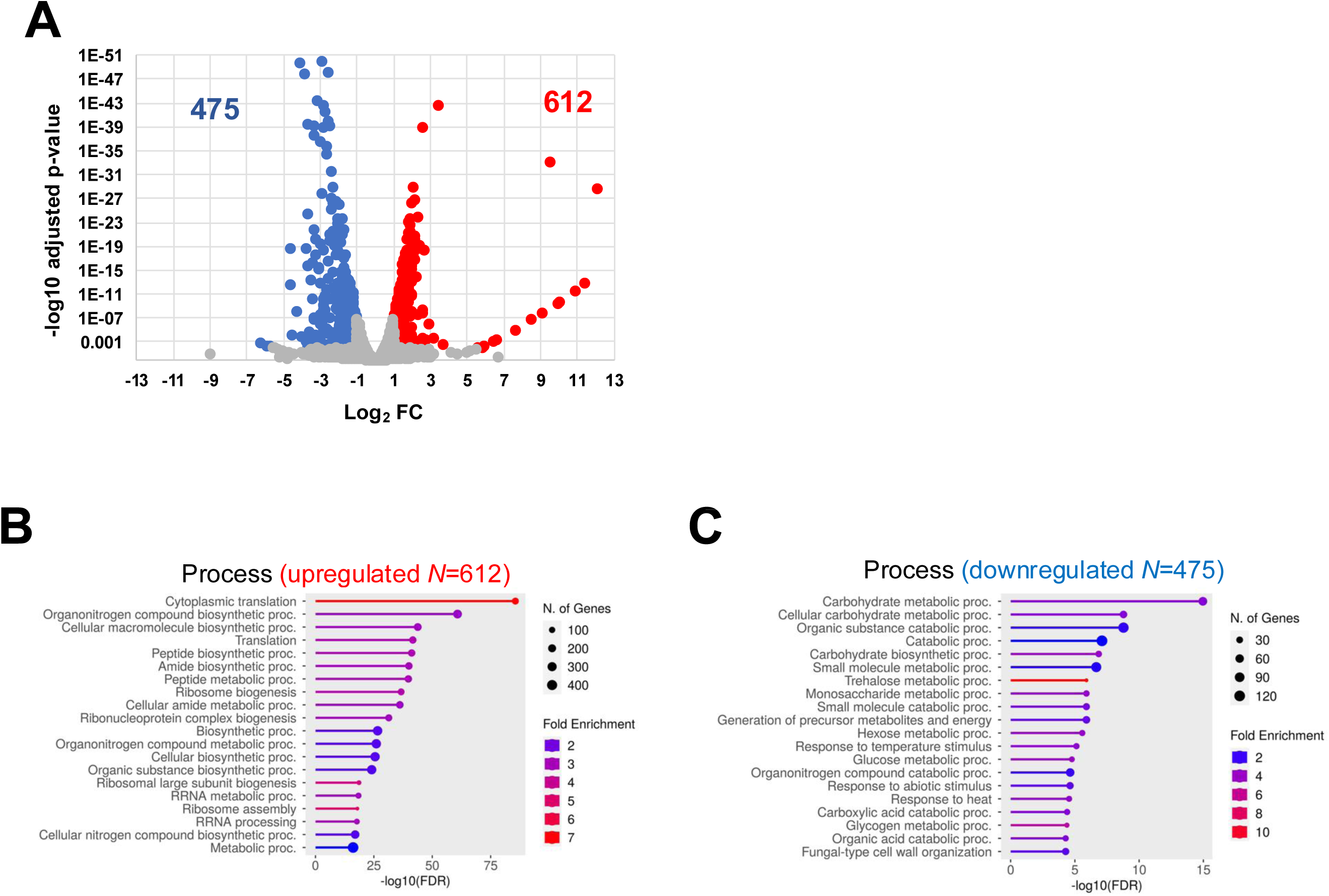
*DEF1* is required for mRNA synthesis and decay. (A). Total RNA-seq was performed in duplicate for both wild-type and *def1Δ* strains. DEseq2 was used to determine changes in expression. *S. pombe* cells were added as spike-in control before RNA isolation to normalize the data. Upregulated transcripts (>2-fold change, p_adj_<0.01) appear in red and downregulated transcripts (<2-fold change, p_adj_<0.01) appear in blue. ShinyGO 0.80 was used to determine the enriched biological processes associated with the upregulated (B) and downregulated (C) genes.

Although a variety of RNA types were sequenced, all differentially expressed transcripts were mRNAs (not shown). We performed Gene Ontology (GO) term analysis on the DEGs; the upregulated genes were associated with ribosomal functions, ribosome biogenesis and translation processes. In contrast, the downregulated transcripts in the mutant were linked to carbohydrate metabolism and various metabolic pathways (Fig. 1B, C, and Supplementary Fig. 1B, C).

Def1 is involved in DNA damage stress response pathways, and cells lacking this gene (*def1Δ* cells) exhibit slow growth (10, 12). To investigate whether changes in gene expression were due to the cells sensing and responding to stress, we compared the differentially expressed genes (DEGs) with the Environmental Stress Response (ESR) genes. ESR genes are regulated by a variety of stressors, including heat shock, DNA damage, nutrient starvation, and oxidative stress, which collectively comprise approximately 900 genes (16). Induced Environmental Stress Response, iESR, genes, or induced are associated with functions such as amino acid transport, redox processes, proteasome activity, and detoxification. In contrast, the Repressed Environmental Stress Response, rESR, group comprises genes that play roles in transcription and translation, as well as ribosomal proteins and those involved in ribosome biogenesis. Our analysis revealed minimal overlap between the upregulated and downregulated differentially expressed genes (DEGs) with the iESR and rESR, respectively (Fig. 2). This finding suggests that the changes in gene expression in *def1Δ* cells are not a result of cellular stress. On the other hand, the opposite was observed. Upregulated DEGs in the *def1Δ* mutant overlapped considerably with the rESR, and the downregulated DEGs overlapped with the iESR (Fig. 2). As mentioned above, the upregulated DEGs are enriched with genes that regulate translation and ribosome biogenesis, and these classes of genes are enriched in the rESR gene set.

**Figure 2.**
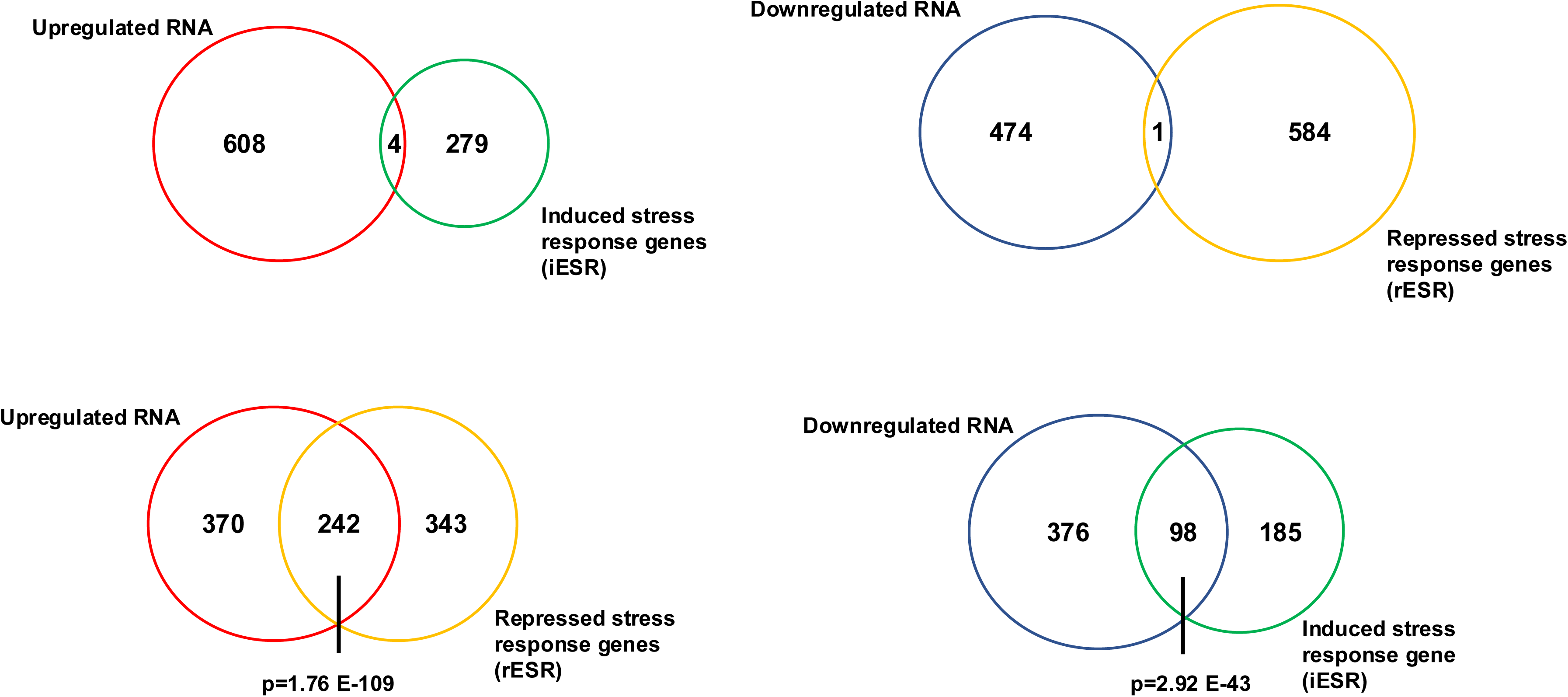
Comparison of gene expression changes to the environmental stress response (ESR) genes. Venn diagrams comparing the overlaps between ESR genes and differentially expressed genes in the *def1Δ* mutant. The collection of induced ESR (iESR) and repressed ESR (rESR) genes (16) were compared to upregulated and downregulated genes in the *def1Δ* mutant strain, as indicated in the figure.

### *DEF1* maintains mRNA synthesis and decay rates

Def1 is recognized for its role in both the initiation and elongation phases of transcription. Therefore, we were surprised to observe so few changes in gene expression, particularly the number with reduced expression levels. RNA sequencing (RNA-seq) measures the steady-state levels of RNA, which is a function of both synthesis and decay. Previous studies have shown that cells can adjust one pathway in response to changes in another to maintain steady-state mRNA levels, a phenomenon known as transcript buffering (1, 4, 17). We therefore investigated the rates of transcript synthesis and decay in the *def1Δ* mutant. We utilized the <u>R</u>NA <u>A</u>pproach <u>T</u>o <u>E</u>quilibrium (RATE-seq) method, which is a pulse-only metabolic labeling technique. In this method, cells are treated with 4-thiouracil for varying labeling durations as the incorporation of the nucleoside analog approaches equilibrium (18). This method accounts for a strain’s doubling time to compensate for slow growing strains. 4-thiouracil -labeled *S. pombe* RNA was used as a spike-in control to normalize between samples and to account for widespread changes in gene expression (see Experimental Procedures section). We conducted RATE-seq on both wild-type and *def1Δ* cells to estimate synthesis and decay rates for 3,938 and 5,631 transcripts, respectively. The data from the replicates showed a very high level of correlation (see Supplementary Fig. 2).

We analyzed the median synthesis and decay rates of 3,596 transcripts that were common to both the wild-type and mutant data sets. In the mutant cells, the global mRNA synthesis rate was significantly reduced. Specifically, the median synthesis rate in the mutant was approximately 3.7-fold lower than that in wild-type cells; however, for some transcripts, the change in synthesis was much more pronounced (see Fig. 3A and Supplementary File 2). Given that the overall synthesis rates were reduced, yet relatively fewer transcripts exhibited lower expression, this suggested that RNA degradation rates were reduced in the mutant. We calculated the decay rates and RNA half-lives and found that, under the experimental conditions, the median decay rate in wild-type cells was 0.259 molecules/min, which corresponds to a median half-life of 2.7 minutes (see Fig. 3B and C and Supplementary File 2). Our estimated half-life is shorter than some previously reported measurements; however, the half-life values found in other studies have varied between 2 and 18 minutes and are highly dependent on the methodology and labeling conditions used (see Supplementary Fig. 3). Most importantly, we found that the decay rate in the mutant was 3.2-fold lower, indicating a global stabilization of mRNAs in these cells (see Fig. 3B and C).

**Figure 3.**
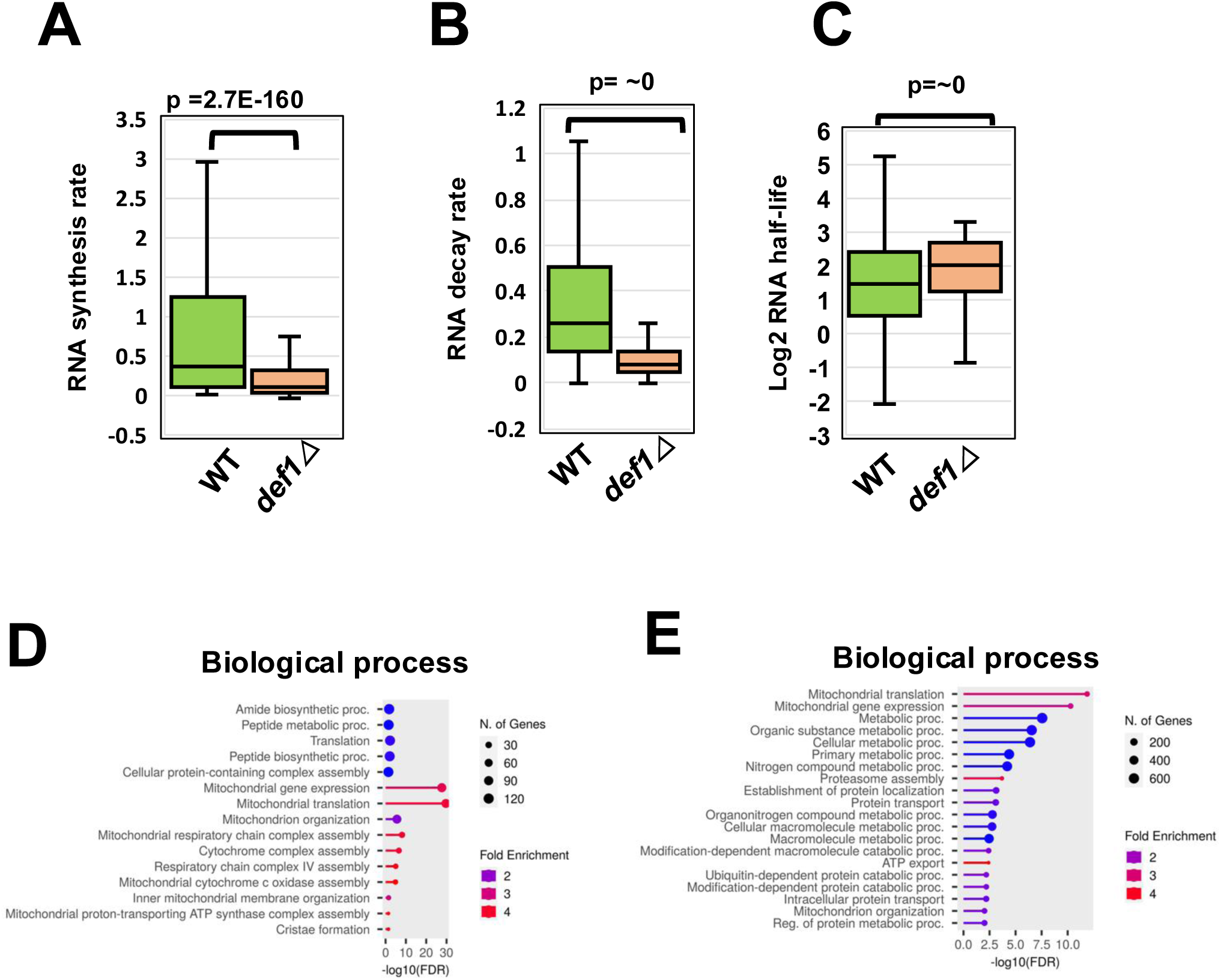
Def1 regulates rates of mRNA synthesis and degradation. (A). RNA synthesis rates are reduced in the *def1Δ* mutant. RATE-seq was used to determine the global rates of RNA synthesis. Rates obtained for the 3596 transcripts that overlap in the wild-type and *def1Δ* samples are displayed in the figure. The p-value was determined using the Wilcoxon rank sum test. Y-axis is molecules/min. (B). Def1 globally controls mRNA decay rates. As in “A”, except decay rates were plotted. (C). As in “A”, except decay rates were used to estimate mRNA half-lives. (D). GO-terms for the biological process of the top 20th percentile of the genes with the highest magnitude of reduced nascent RNA synthesis (*def1Δ*/wild type). GO terms were determined using ShinyGO 0.80. (E). GO-terms for the biological processes of the top 20th percentile of mRNAs with the largest change in mRNA decay.

If Def1 regulates the RNAs of genes involved in specific cellular processes or pathways, we anticipate that its effects on mRNA metabolism will be most significant for those transcripts. To investigate whether Def1 targets mRNAs within a particular pathway, we performed Gene Ontology (GO) term analysis on the transcripts in the top 20th percentile for synthesis (Fig. 3D) and half-life fold change (Fig. 3E). The biological processes most affected by the deletion of *DEF1* included mitochondrial translation, protein metabolism, and translation.

We then plotted the changes in decay rates of individual mRNAs against their changes in synthesis rates (Fig. 4). We observed a strong positive correlation (R = 0.834), which indicates that transcript buffering is occurring in the mutant. This suggests that Def1 is not necessary for transcript buffering, and the corresponding changes in synthesis and decay rates explain why no global (widespread) change in steady-state mRNA levels was observed in the mutant.

**Figure 4.**
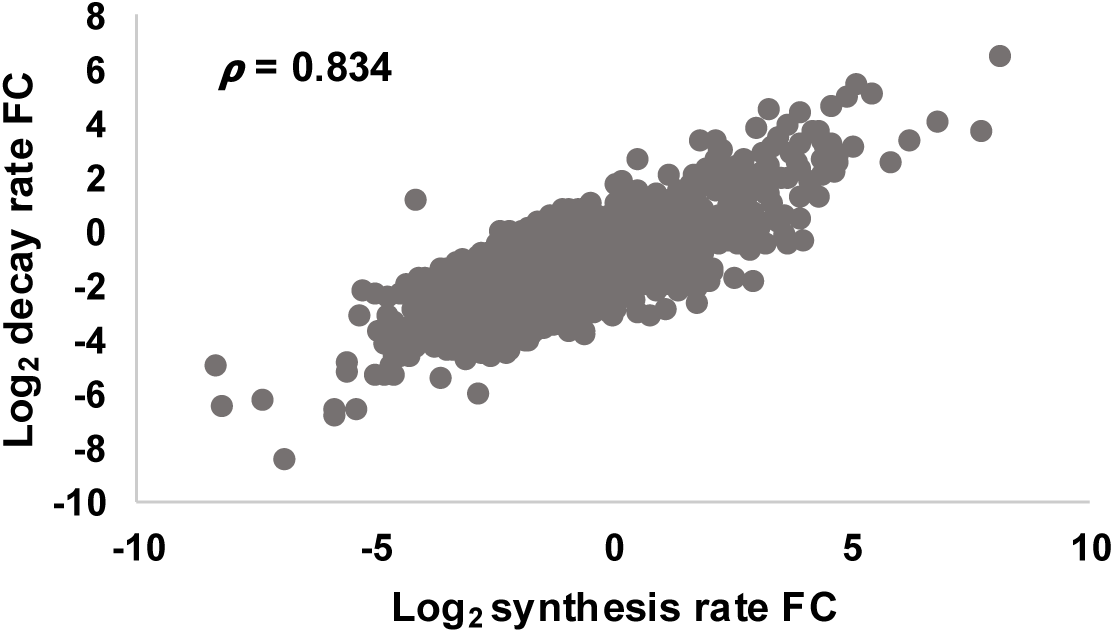
The *def1Δ* mutant displays gene expression buffering. A scatterplot was constructed by plotting the fold change (FC) in RNA decay and synthesis rates plotted on the y- and x-axis, respectively. The Spearman correlation coefficient is shown in the scatterplot.

### Def1 interacts with cytoplasmic mRNA regulatory proteins

Proximity labeling, using an engineered biotin ligase (BioID), is a technique employed to identify protein-protein interactions within cells, allowing for the detection of transient and weak interactions (19, 20). Recently, we conducted BioID experiments on subunits of the Ccr4-Not complex and discovered that several bait proteins strongly labeled Def1 (14). Additionally, the human homolog of Def1, UBAP2/2L, was found to interact with hCcr4-Not subunits (15). Therefore, we performed a reciprocal experiment by fusing the TurboID (TID) version of the biotin ligase enzyme to the C-terminus of Def1 (21). Deleting *DEF1* causes slow growth and increased sensitivity to DNA-damaging agents (10, 12). We verified that the fusion of TID to the C-terminus of Def1 did not affect cell growth, even in the presence of hydroxyurea, suggesting that the addition of TID to Def1 is well tolerated. (Supplementary Fig. 4A).

We expressed “free” TID at levels close to those of Def1-TID as a control (Supplementary Fig 4B,C). The interacting proteins were identified by calculating the fold enrichment of Def1-TID labeled proteins versus free TID. Labeled proteins were subjected to mass spectrometry, and we implemented a cutoff of 1.5-fold enrichment and p < 0.05 to identify interacting or proximal partners of Def1 (Fig. 5A and Supplementary File 3). A total of 97 proteins met these criteria, with 42 involved in various aspects of RNA regulation. Notably, Def1-TID also labeled retrotransposon proteins (Ty), which were not included in the 97 (Supplementary File 3). The significance of this is not known. We submitted the enriched proteins to Gene Ontology (GO) term analysis, which returned terms associated with the regulation of translation, post-transcriptional control, and mRNA decay. The analysis for cellular component yielded terms such as ribonucleoprotein granule, P-body, translation initiation factor 4F, polysome, and the Ccr4-Not core complex. (Fig. 5 B-D). Overall, Def1-TID predominantly labeled cytoplasmic RNA regulators.

**Figure 5.**
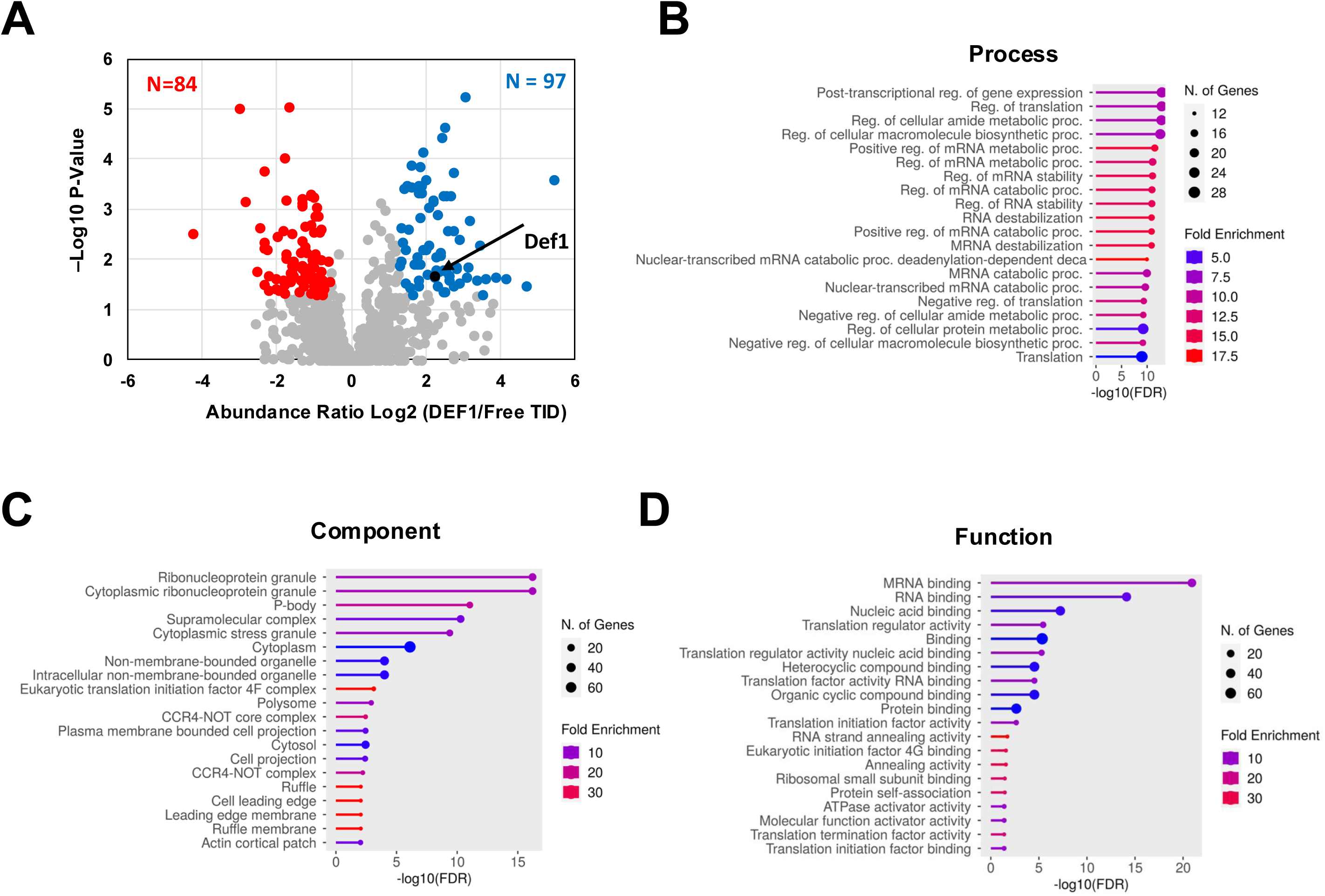
BioID analysis reveals that Def1 interacts with cytoplasmic posttranscriptional regulatory factors. (A). BioID labeling was performed in cells expressing Def1-Turbo ID (TID) and cells expressing free TID enzyme. Biological replicates were analyzed. Biotinylated proteins pulled down by streptavidin beads were identified by mass spectrometry. Data are presented as a Volcano plot of the fold change in protein abundance ratio of Def1-TID versus free TID (Log2) versus -log10 p-value. The enriched proteins (>1.5 fold, p<0.05) appear in blue and underrepresented proteins (>-1.5 fold, p<0.05) appear in red. Proteins enriched in the Def1-TID sample were analyzed using ShinyGO 0.80, and biological process (B), component (C), and function (D) are shown.

Eighty-four proteins were found to be underrepresented in the Def1-TID sample compared to the free TID control sample. Previous BioID studies have shown that proteins can be underrepresented in the test sample relative to the free control (14). This underrepresentation may be due to the fusion of the test protein to TID, which might prevent the TID enzyme from labeling non-targets non-specifically within the cell. When we subjected the underrepresented proteins to Gene Ontology (GO) term analysis, we found broad terms with weak false discovery rates (FDR) and enrichment scores (not shown), suggesting a lack of biological significance.

It was surprising that Def1 did not label many RNAPII transcription factors or other nuclear proteins involved in mRNA synthesis. The predominant localization of Def1 in the cytoplasm, particularly in the absence of transcription stress, likely explains why few nuclear proteins were labeled. Additionally, the biotin ligase was fused to the C-terminus of Def1, and the C-terminus retains Def1 in the cytoplasm (12). These data suggest that Def1 interacts with cytoplasmic RNA-regulating proteins, indicating that it may have post-transcriptional functions, such as mRNA storage, translation, or decay.

### Tethering Def1 directly to an mRNA reduces its expression and accelerates its decay

The extended half-lives of mRNA in the *def1Δ* mutant, along with the labeling of various mRNA deadenylation and decapping factors by Def1-TID, suggest that Def1 regulates mRNAs in the cytoplasm. However, the decreased mRNA turnover observed could be a compensatory response to a lower synthesis rate caused by transcript buffering (1, 4). To provide evidence that Def1 regulates mRNA turnover, we employed an MCP/MS2 tethered reporter assay, which has been previously used to demonstrate the direct regulation of mRNA by decay factors (22). The reporter construct produces an mRNA that includes an open reading frame (ORF) for green fluorescent protein (GFP) along with MS2 stem loops in the 3’ UTR (Fig. 6A). We fused MCP to the N-terminus of Def1 (labeled as MCP-Def1). The MCP-Def1 fusion protein complemented a *def1Δ* strain, demonstrating its functionality (Supplementary Fig. 5A). The expression levels of the MCP-Def1 fusion protein were similar to those of the free MCP protein (Supplementary Fig. 5B and C). As a positive control, we tested a Dhh1-MCP fusion protein, which has been previously shown to repress reporter mRNA (22, 23). We measured the levels of GFP protein produced by the reporter. Tethering Def1 to the mRNA resulted in a significant decrease in GFP protein levels compared to the free MCP control (Fig. 6B). Specifically, GFP protein levels were reduced to approximately 50% in cells expressing the Def1 fusion protein, although this reduction was not as pronounced as the repression observed with the Dhh1-MCP control. Next, we assessed mRNA levels using quantitative PCR (qPCR). The amount of GFP mRNA was normalized to that of *ACT1* and plotted (Fig. 6C). The results showed that the mRNA levels of the reporter were lower in cells expressing MCP-Def1 compared to those expressing MCP alone. Furthermore, Dhh1-MCP was slightly more effective at repressing the reporter mRNA expression.

**Figure 6.**
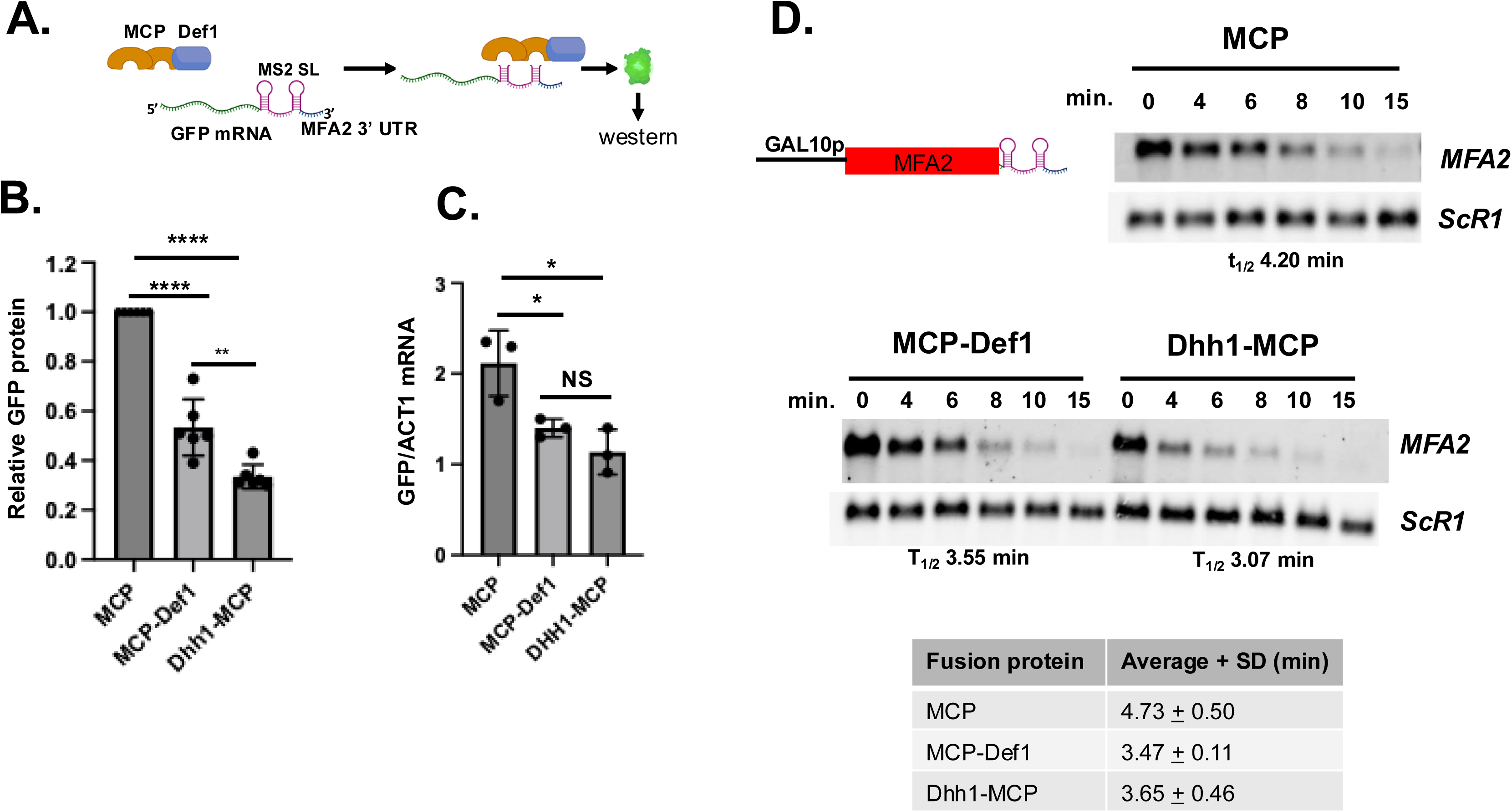
MS2-MCP tethering reporter assays suggest that Def1 functions in posttranscriptional control of mRNAs. (A) Schematic of the MS2 coat protein (MCP)-MS2 tethering assay. MCP-Def1 has MCP fused to the N-terminus; Dhh1-MCP was used as a positive control. Expression of the free MCP and the MCP-Def1 fusion proteins was driven by the *DEF1* promoter. Dhh1-MCP was driven by the *DHH1*-promoter. (B) The relative abundance of GFP protein in the fusion protein-expressing cells, compared to that expressing free MCP protein, was set at 1.0. Taf14 was used as a loading control, and GFP protein signals were normalized to the loading control. The averages of three biological replicates from two experiments are shown (N=6). The error bars represent the standard deviations. P values were calculated using an unpaired t-test. ****, p<0.0001; **, p<0.01. (C). Graph of steady-state GFP mRNA. RT-qPCR measured the abundance of both GFP and *ACT1* (control) mRNA. GFP mRNA amounts were normalized to *ACT1*. Biological triplicates were analyzed (N=3). P values were calculated using an unpaired t-test. *. P<0.05. (D, upper). Cells transformed with a GAL10p-MFA2-MS2 reporter gene (22) and each of the MCP derivatives described in the panel were grown in galactose-containing dropout medium, transferred to dextrose-containing medium and samples were collected at various time points following transcription shutdown. The levels of *MFA2* RNA were detected by Northern blotting. *ScR1* was the loading control. The calculated half-lives, after correcting for loading, are listed below the panel. (D, lower) *MFA2* reporter mRNA half-lives from multiple determinations (N = 4). The signals from *MFA2* were normalized to those of the loading control *ScR1*. The averages and standard deviations of reporter mRNA half-lives are reported. P-values were calculated using a two-tailed unpaired t-test. The p-values comparing the half-life between cells expressing MCP and MCP-Def1 and Dhh1-MCP were p < 0.01 and p < 0.054, respectively.

We investigated whether tethering Def1 to an mRNA could reduce the half-life of an RNA reporter. To achieve this, we used a reporter construct that included the MFA2 open reading frame (ORF) followed by a 3’ untranslated region (UTR) containing MS2 binding sites, all under the control of the *GAL10* promoter. We shut down transcription by shifting the cells from a medium containing galactose to one with dextrose. Figure 6D presents a representative experiment and the average results from multiple experiments (N=4). In cells expressing only the MCP protein, the reporter mRNA had an average half-life of approximately 4.2 minutes. Conversely, cells expressing the MCP-Def1 fusion protein showed a significantly greater enhancement in decay, reducing the average half-life to about 3.6 minutes (p < 0.01). The reduction in half-life observed with the MCP-Def1 fusion was comparable to the Dhh1 control, which had an average half-life of around 3.7 minutes (p < 0.054). These reporter assays provide evidence that Def1 plays a role in regulating mRNAs post-transcriptionally.

Def1 functions similarly to other factors involved in both the decay and synthesis of mRNAs. Previous studies have demonstrated that tethering mRNA decay factors or general transcription factors to promoters can activate transcription (2, 24). Def1 copurified with the general transcription factor TFIIH, was crosslinked to promoters of genes and affected start site selection in an *in vitro* transcription assay, suggesting it may regulate initiation (13). Def1 has been shown to copurify with the general transcription factor TFIIH and influence start site selection in *in vitro* transcription assays (13). Given this information, we tested whether recruiting Def1 to a promoter would enhance transcription. A reporter gene containing β-galactosidase under the control of 8 LexA operators was used. We constructed fusion proteins combining the DNA-binding domain of LexA with full-length Def1. The expression of both LexA alone and the LexA-Def1 fusion was driven by the *DEF1* promoter, as described in the methods section. For comparison, we also analyzed the LexA-TBP (TATA-binding protein) fusion as a positive control. While the LexA-TBP fusion strongly activated the reporter gene, the LexA-Def1 protein did not (see Supplementary Fig. 6). However, it is important to note that this tethering assay assesses the effects on preinitiation complex formation, and a negative result does not rule out Def1’s potential impact on transcription, particularly during the elongation phase.

## Discussion

An increasingly large number of gene regulatory proteins are being discovered to have new or previously overlooked functions. For instance, Ccr4-Not, originally believed to regulate TATA-binding protein (TBP) and transcription initiation primarily, is now acknowledged for its significant role in the post-transcriptional control of mRNAs. Additionally, Degradation Factor 1 (Def1) has been identified as a key player in mitigating DNA damage. Extensive research has established Def1 as a factor that facilitates the elongation process and degrades Rpb1, thereby clearing stalled RNA polymerase II (RNAPII) from genes. Other nuclear functions have also been attributed to Def1 (for review, see (6)). Excluding Def1 from the nuclear compartment is crucial to prevent interference with transcription (as noted in (12)). Although Def1 is present in the cytoplasm, its cytoplasmic functions have not been identified. In this study, we report a new role for Def1 in the cytoplasm related to post-transcriptional gene control.

Def1 has long been recognized as an RNA polymerase II (RNAPII) elongation factor, co-purifying with TFIIH and influencing start site selection in reconstituted transcription reactions (6, 12, 13). Our study describes the first transcriptome analysis of this transcription factor. Consistent with its expected role in transcription, the deletion of *DEF1* globally reduced mRNA synthesis (Fig. 3A). Its function in maintaining mRNA decay rates was unexpected, suggesting a role in the cytoplasm. While the reduced synthesis rates in the mutant may contribute to the slowed mRNA decay due to transcript buffering, we present additional evidence that Def1 plays a direct role in mRNA decay. First, Def1 binds many cytoplasmic post-transcriptional regulators, including the major deadenylase complex, Ccr4-Not, as demonstrated by BioID labeling. Both Def1 and Ccr4-Not are involved in regulating transcription elongation and the degradation of Rpb1 during DNA damage stress, which could have explained their interaction (6, 12, 25). However, evidence for a function in decay comes also from the MS2/MCP tethering reporter assays. The recruitment of Def1 to an mRNA accelerated its decay and repressed its expression. Moreover, proximity labeling experiments using Def1, Not1, and Not4 as baits revealed reciprocal interactions in cells (Fig. 5 and (14)). This suggests that Def1’s ability to control mRNA decay may be mediated by recruiting decay factors, such as Ccr4-Not, to mRNAs. We attempted to examine this by conducting the MS2/MCP reporter assay in a *ccr4Δ* mutant. However, we found that the expression of Def1-MCP protein in the *ccr4Δ* mutant was severely compromised, reduced to only 20% of the levels observed in wild-type cells (not shown), which made the assay results difficult to interpret. Additionally, Def1 also labeled the decapping enzyme Dcp2, indicating that Def1 could influence this step in mRNA degradation as well. Further studies are needed to clarify how Def1 regulates these processes. The recruitment of decay factors may involve the polyglutamine-rich C-terminus of the protein, as this is the location where the TID enzyme was fused.

A cryptic nucleic acid-binding domain has been identified in Def1, specifically between positions 1-207 (26). This domain may bind to transcripts non-specifically; however, it is more probable that Def1 is recruited to mRNAs via the Ccr4-Not complex or through sequence-specific RNA-binding proteins. The Def1-TID protein biotinylated multiple sequence-specific RNA-binding proteins, including all members of the Pumilio family of RNA binding proteins (Puf1/Jsn1, Puf2, Puf3, and Puf4) (Supplemental File 3). These proteins bind to specific sequences in the 3’ untranslated regions (UTRs) of RNAs, marking them for degradation and translational repression (27). Interestingly, Puf3 is involved in targeting and promoting the degradation of mRNAs of nuclear-encoded mitochondrial proteins (28, 29), and the mRNAs that are most sensitive to the loss of Def1 encode for mitochondrial proteins (Fig. 3D and 3E).

Def1 interacts with ribonucleoprotein granule (RNP) markers, including Ccr4-Not subunits, Edc3, Dhh1, and Lsm7. Ribonucleoprotein granules, such as stress granules and processing bodies, are cytosolic compartments that facilitate translational repression, where mRNAs are stored alongside initiation and decay factors during times of stress (30). However, neither UV stress (12) nor nutrient stress (data not shown) induced the formation of Def1-containing foci, such as processing bodies. Recently, UBAP2/2L was identified as the putative human homolog of Def1, as it has been shown to direct the ubiquitylation of the large subunit of RNA polymerase II, Rpb1(31). UBAP2/2L is located in stress granules and is necessary for their formation (32, 33). While UBAP2/2L possesses a ubiquitin-binding domain similar to that of Def1, it lacks the polyQ-rich domain found in Def1. Therefore, it was unclear whether Def1 has cytoplasmic functions or if UBAP2/2L acquired these functions. Although Def1 does not form detectable cytoplasmic foci, it may regulate the partitioning of mRNAs between the repressed pool and the actively translated pool. Not4 and Not5, members of the Ccr4-Not complex, play a crucial role in this partitioning, determining the distribution of mRNAs between the soluble and insoluble fractions (34).

Our research addressed the perplexing cytoplasmic localization and the remarkably high abundance of Def1. According to the Saccharomyces cerevisiae database (SGD), the median abundance of Def1 is estimated to be around 21,000, but some estimates predict the copy number as high as 80,000 per cell. As mentioned earlier, DNA damage leads to the proteasomal processing of a portion of Def1, while a larger fraction of the unprocessed protein remains in the cytoplasm. Def1 has functions in the cytoplasm, and its presence in this part of the cell serves a purpose beyond merely preventing interference with transcription in the nucleus, as previously suggested (12). Def1 contributes to both transcription and crucial cytoplasmic functions, thereby adding it to the list of factors involved in transcription within the nucleus, as well as mRNA decay in the cytoplasm. Therefore, Def1 may play a vital role in nuclear-cytoplasmic communication by coordinating transcription and decay.

## Experimental Procedures

### Yeast strains and plasmids

Strains and plasmids used in this work are contained in Supplementary Tables 1 and 2, respectively. Cells were grown in rich media, YPAD (2% Bacto-Peptone, 1% yeast extract, 20 μg/ml adenine sulfate, 2% dextrose) or synthetic dropout media made with yeast nitrogen base plus ammonium sulfate and either 2% galactose or dextrose. Gene deletions were produced by homologous recombination using PCR-generated cassettes (35). The genotypes of the mutants were screened by PCR of genomic DNA. Plasmids were sequenced. Details on the construction are available upon request.

### Growth spot assay

Yeast strains were grown to saturation at 30°C in 5 mls of the appropriate medium. Serial dilutions were prepared in sterile distilled water, and diluted samples were spotted on plates containing the appropriate media and supplements. Plates were incubated at 30°C for 24 hr or more, as indicated in the respective figures.

### Western blotting

Samples were separated on either SDS-PAGE or Tris-acetate gradient gels, and proteins were transferred to nitrocellulose membranes by semi-dry transfer. Membranes were stained with ponceau S, imaged, and then de-stained. Membranes were blocked in 5% milk in TBST 50 mM Tris-HCl, pH 7.5, 0.15 M NaCl, 0.1 mM EDTA, and 0.1% Tween-20 for at least 1 hour and then probed with the primary antibody in 2% or 4% milk for at least 2 hr. Antibodies to Taf14 and Dhh1 have been described in other publications and are custom antibodies raised in rabbits, validated by analyzing an extract from cells with the gene deleted (14, 36). Antibodies to Def1 were raised in rabbits to a 6HIS-Def1 (1-500) fusion. The Def1 was validated by probing an extract from a *def1Δ* strain. Commercial primary antibodies used are Anti-FLAG M2 (Millipore-Sigma), GFP (Y1030,UBP-Bio) and anti-HA, (HA.11, Biolegend). Epitope tag antibodies were validated by including a sample from untagged cells and observing mobility changes in proteins of different sizes. Signals were detected by fluorescence or chemiluminescence (ECL) and imaged on X-ray film or a ChemiDoc MP imaging system (Bio-Rad). Statistical analysis and graph construction were performed in Prism GraphPad.

### BioID

Turbo ID (TID) was fused to the C-terminus of Def1, and truncation derivatives, using PCR-derived cassettes from pJR106 as described in (14). Free TID was expressed from the *CHA1* promoter, and this strain was used as the control. Protein extraction and isolation on streptavidin magnetic beads (Pierce, Thermo-Fisher, #88817) was conducted as described in (14). Biological duplicates were analyzed. Proteins were digested on the beads with 1 ug of trypsin (Promega Trypsin Gold) after alkylation with iodoacetamide overnight at 37°C. Purified peptides were labeled with TMTpro 16plex reagents (Thermo-Fisher, #A44520) per the manufacturer’s recommended protocol. Peptides were analyzed on a Nano-flow Liquid Chromatography (Thermo Easy-nLC 1200). The peptides were loaded on an Acclaim PepMap100 trapping column (75 mm × 2 cm, C18, 5 mm, 100 Å, Thermo), and separated on an Acclaim PepMap RSLC column (50 mm × 15 cm, C18, 2 mm, 100 Å, Thermo) at a flow rate of 300 nL/min in the following gradient of mobile phase B (80% ACN in 0.1% aqueous FA): 5% - 50% B in 210 min, 50% - 90% B in 30 min. A Thermo Orbitrap Eclipse mass spectrometer was operated in a data dependent mode using a method based on the TMT MS2 template. The resulting mass spectra were processed using Thermo Proteome Discoverer 3.1 software.

### Total RNA-seq for steady state mRNA levels

Cells were grown in YPAD and harvested at log phase (OD600∼0.8-1). Cells were washed with nuclease-free water and resuspended in AE-buffer (50 mM sodium acetate pH 5.3, 10 mM EDTA,1% SDS). Cells were spiked with *S. pombe* cells at a ratio of 14:1. RNA was isolated using AE-phenol chloroform extraction, followed by overnight ethanol precipitation with sodium acetate. RNA samples were washed with 80% ethanol and resuspended in nuclease-free water. RNA was quantified on Nanodrop, and the quality was verified by analysis on a Tapestation. RNA samples with RIN scores >9.0 were used. Biological replicates were analyzed.

### RNA metabolic labeling studies

The procedure is a derivative of the one used in a previous publication (37). *Saccharomyces cerevisiae* cells growing in YPD were treated at log phase (OD_600_∼0.8-1) for 6-, 9-, 12-, 24- and 90-min with 0.65 mg/ml 4-thio-uracil (Millipore-Sigma). Thirty OD_600_ cultures were transferred to 2 volumes ice-cold methanol on dry ice to halt 4tU incorporation. Cells were harvested by centrifugation at 3000 rpm for 3 min, and resuspended in AE-buffer (50 mM sodium acetate pH 5.3, 10 mM EDTA,1% SDS). The *S*. *cerevisiae* cells were spiked with pre-labeled *S. pombe* cells at a ratio of 1:14. *S. pombe* spike-in was generated by growing cells in YPD to log phase (OD_600_∼0.8) and treated with 5 mM 4tU for 12 min. RNA was isolated using AE-phenol, then extracted with chloroform, and finally precipitated overnight in ethanol. RNA samples were resuspended in nuclease-free water. RNA was subjected to biotinylation (120 μg RNA, 20 mM HEPES, pH 7.4, 12 μg MTSEA biotin-XX (Biotium Co.) in a final volume of 600 μL for 30 min at room temperature). Free MTS-biotin reagent was removed by phenol-chloroform-isoamyl alcohol extraction and precipitation with NaCl and isopropanol. Dynabead MyOne streptavidin C1 beads (Thermo-Fisher) were washed twice in high salt wash buffer, HSWB, (100 mM Tris-HCl pH 7.4, 10 mM EDTA, 1M NaCl) and blocked with blocking buffer (100 mM Tris-HCl ph 7.4, 10 mM EDTA, 1M NaCl, 40 ng/ul glycogen), then resuspended in HSWB. 80 ug of biotinylated RNA was incubated at 65°C for 7 min and streptavidin pull down was performed using 100 ul blocked beads and then washed thrice with HSWB, and eluted twice in 25 ul elution buffer (100 mM DTT, 10 mM EDTA, 1M NaCl, 0.05% Tween-20). The RNA was concentrated and quantified on a Qubit instrument (Life Technologies).

### Library preparation and NextGen sequencing

Total RNA was depleted of rRNA using the Qiagen FastSelect (#334215) kit. Libraries for steady-state RNA and RATE-seq samples were prepared using the Illumina Total Stranded library prep kit and the Illumina UDI indexes (#20040529). The libraries were analyzed on TapeStation-4150 at the Genomics Core Facility, Penn State University, and were quantified using NEBNext® Library Quant Kit for Illumina®. Libraries were sequenced on the Illumina NextSeq 2000. Illumina P2 100 cycle reagent cartridge (#EC1591461-EC11) was used for paired-end sequencing. Reads were mapped to the sacCer3 genome using the ENCODE 3 pipeline. Spike-in reads were mapped to the *S. pombe* genome (https://www.encodeproject.org/pipelines/ENCPL002LPE/). Star was used to map the reads, and RSEM-1.2.28 to get estimated counts and TPMs (Genes: Saccharomyces_cerevisiae.R64-1-1.106.ucsc.gtf). DESeq2 was used to determine the differential expression of genes (38). The truncated expected counts from the RSEM output were used as input for DESeq2. Genes with <15 reads across samples were filtered out. The *p_adj_* value of 0.01 was used as a threshold.

### RATE-seq data modeling

RNA reads obtained from Deseq2 were used for nonlinear regression modelling to obtain decay rates in *R* using the *nls.table* function. The model for RNA decay is (18):

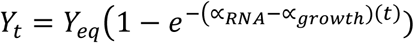

Where

Y_t_ = Amount of labelled transcript at time *t*

Y_eq_ = Abundance of labelled transcript at steady state

t = labeling time

RNA decay rate constant, ∝= (∝*_RNA_*−∝*_growth_*)

α_RNA_ = RNA decay rate

α_growth_ = Growth rate = *ln2/doubling time* = 0.0077 (WT) or 0.0046 (*def1Δ*)

Synthesis rate was estimated in Excel from the decay rate constant using the formula:

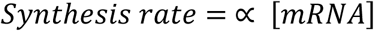

Where mRNA abundance is estimated from steady state read counts, assuming a total of 60000 mRNA molecules per cell(29).

mRNA half-life was estimated in Excel from decay rate constant using the formula:

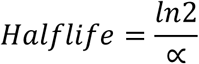

### mRNA decay and transcription reporter assays

The MS2-MCP mRNA reporter assay was conducted as described in a previous publication (22). Plasmids expressing MCP fusions with Def1, were constructed from gene synthesis and PCR-generated fragments using inPhusion assembly (Takara Bio) (Supplemental Table 2). Fusion proteins were driven by the *DEF1* promoter and contained its 3’UTR. MCP fusion protein expression plasmids were co-transformed with the reporter vector pJC428 or pJC2 into W303 cells (Supplemental Table 1). Cells were grown in triplicate in -uracil/-leucine dropout media to an OD_600_ of 1.0. The expression levels of the MCP fusion proteins were verified using anti-FLAG antibodies, and the levels of the reporter GFP protein using anti-GFP antibodies. To calculate the levels of GFP protein, a serial dilution standard curve of an extract was included on the blots. Loading was normalized to Taf14 levels, which served as a loading control. Signals from images were quantified using ImageJ. An unpaired t-test was used to calculate the p-values. Steady state RNA levels of GFP and *ACT1* were quantified by qPCR using primers described in (22). Statistical analysis and graph construction were performed in Prism GraphPad.

mRNA half-life measurements were conducted using a GAL10p-MFA2-MS2 reporter plasmid (22). To measure the half-life of the reporter mRNA, cells were grown to mid-log phase in dropout media containing 2% galactose and then collected by centrifugation. After resuspension in dropout media lacking a carbon source, an aliquot was removed and quenched with a half-volume of methanol chilled to -70 °C. Dextrose was then added to 4% and aliquots were removed at different time points and quenched in methanol chill to -60°C. RNA was subjected to Northern blotting using random-primed biotinylated DNA probes (BioPrime Kit,Thermo-Fisher) to detect *MFA2*. After washing, RNA was detected using streptavidin-IR800 (926-32230, Li-Cor) and scanned on a Chemidoc-MP (Bio-Rad). Blots were stripped of the *MFA2* probe and then probed with *ScR1* as a loading control.

The Beta-galactosidase transcription reporter assay was conducted by co-transforming W303 cells with pSH18-34 and plasmids expressing LexA fusion proteins with Def1 (Supplementary Table 2). Fusion proteins were driven by the DEF1 promoter and contained its 3’UTR. Cells were grown in triplicate in -uracil/-leucine dropout media containing 2% dextrose to an OD_600_ of 1.0. Protein extraction and beta-galactosidase assays were performed as described in a previous publication,(39).

### RT-qPCR

RNA samples were quantified on Nanodrop and run on 1% agarose gel to confirm the quality. 1 ug of RNA was treated with 1U DNaseI for 30 min at 37°C followed by quenching with 5 mM EDTA at 65°C for 10 min. RevertAid RT kit (Thermo-Fisher,FERK1622) was used for cDNA preparation according to the manufacturer’s protocol using a mix of random and oligo dT primers. Serial dilutions of cDNA were used to set up qPCR reactions. PCR reactions were performed with PerfeCTa SYBR green SuperMix and using an Agilent Real-Time machine. The data was analyzed using the Agilent AriaMx software and Excel.

## Supporting Information

This article contains supporting information

## Data availability

Genomics data have been deposited to GEO under accession number GSE290861. Proteomics data were uploaded to MassIVE under MSV000097607.

## Acknowledgements

Mass Spectrometry was performed by Dr. Tatiana Laremore at the Penn State Huck Proteomics and Mass Spectrometry Core Facility (RRID:SCR_024462). The Genomics Research Incubator (RRID:SCR_024530) and Genomics (RRID:SCR_023645) cores of the Huck Institutes of Life Science were used in this work. Karen Arndt is recognized for sharing the nascent transcription protocol. Jeff Coller is thanked for providing the plasmids used in the MS2/MCP reporter assays. This research was supported by funds from the National Institutes of Health (R35 GM136353 to J.C.R).

## Author contribution Statement

O.T.A., M.M.H. and J.C.R. conducted wet bench experiments. O.T.A., S.K., and B.M.G provided bioinformatics support and analysis. C.A.K. assisted with library construction and Illumina sequencing. O.T.A. and J.C.R. prepared the figures. O.T.A. and J.C.R. wrote the paper.

## Conflicts of Interest

## Abbreviations

GO: Gene Ontology
ESR: Environmental Stress Response
RATE-seq: RNA Approach To Equilibrium
TID: Turbo ID biotin ligase
FC: fold-change
DEG: differentially expressed genes
4tU: 4-thio-uracil
UTR: untranslated region
MCP: MS2 coat protein

