## supplemental figures for "Gene expression analysis and proximity labeling reveal posttranscriptional functions of the yeast RNA Polymerase II regulator Def1"

### Supplementary Figure 1

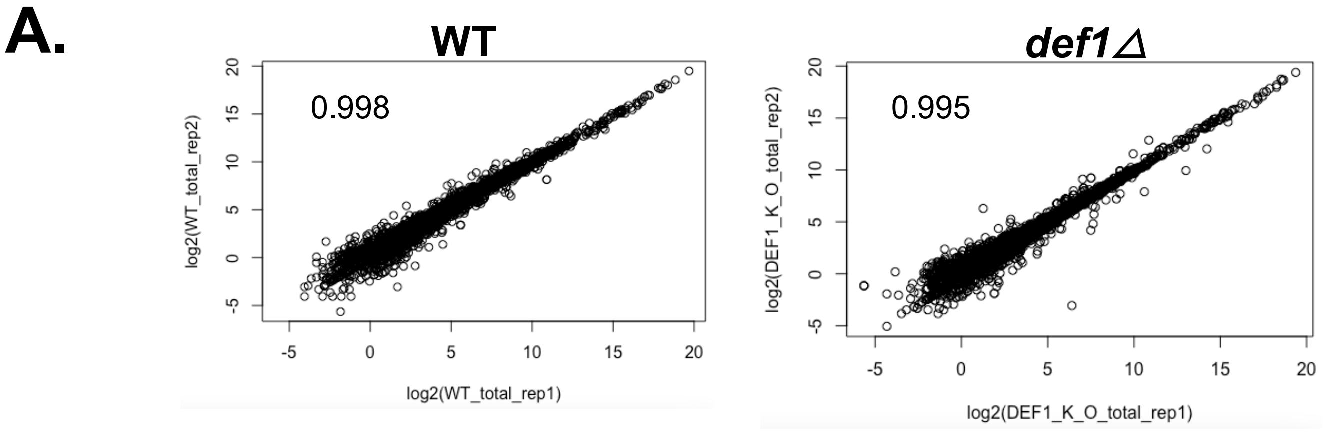


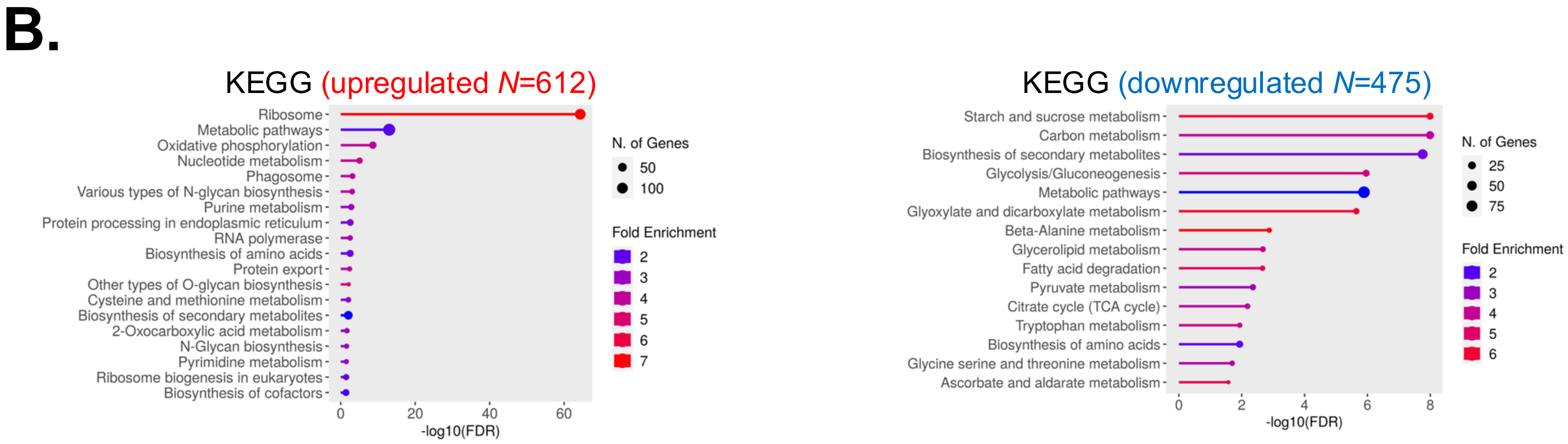

#### Supplementary Figure 2

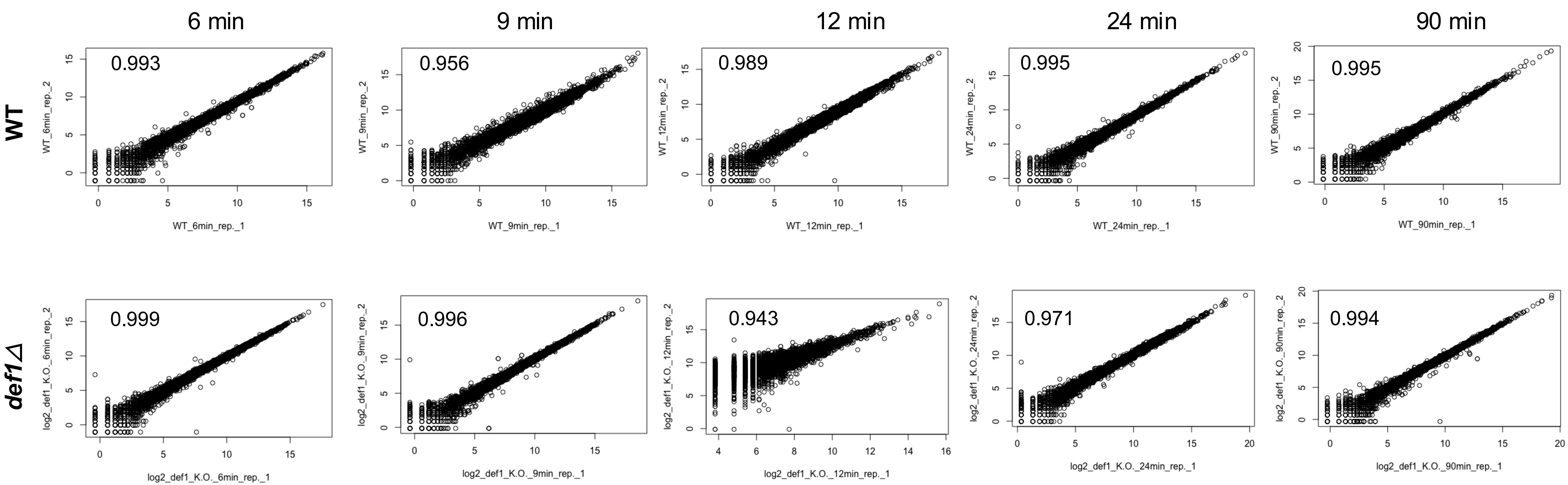

### Supplementary Figure 3

| **Article** | **Approach** | **Median halflife** |
|---|---|---|
| Miller *et al*. | DTA (4sU, 5mM) | 11 min |
| Munchel *et al*. | 4TU pulse-chase (4TU, 0.2mM) | 18 min |
| Neymotin *et al*. | RATE-seq (4TU, 0.5mM) | 10 min |
| Presynak *et al*. | RNAPII inactivation | 7.4 min |
| Chan *et al*. | 4-TU pulse only (4TU, 1mM) | 3.6 min |
| Alalam *et al*. | SLAM-seq (4TU, 0.2mM) | 9.4 min |
| Baudrimont et al. | Promoter shutdown | 2 min |
| This study | RATE-seq (4TU, 5 mM) | 2.7 min |

#### Supplementary Figure 4

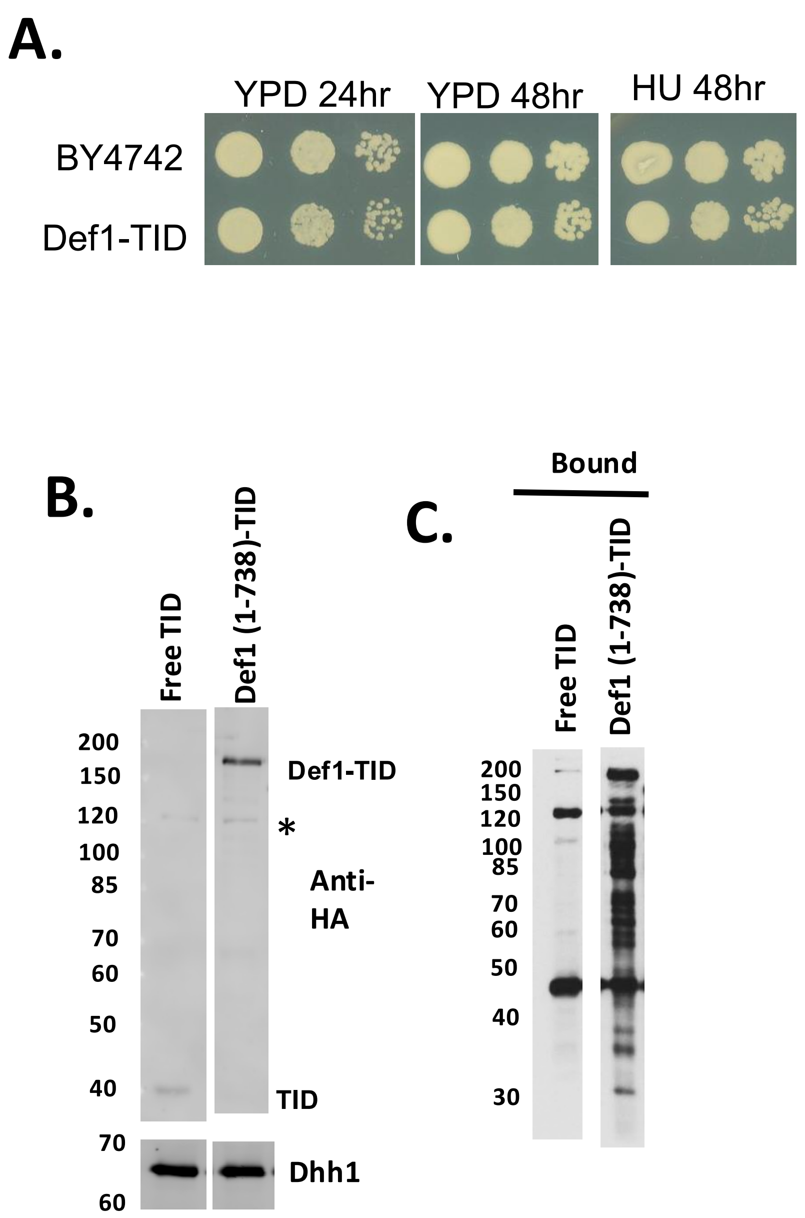

Supplementary figure 5

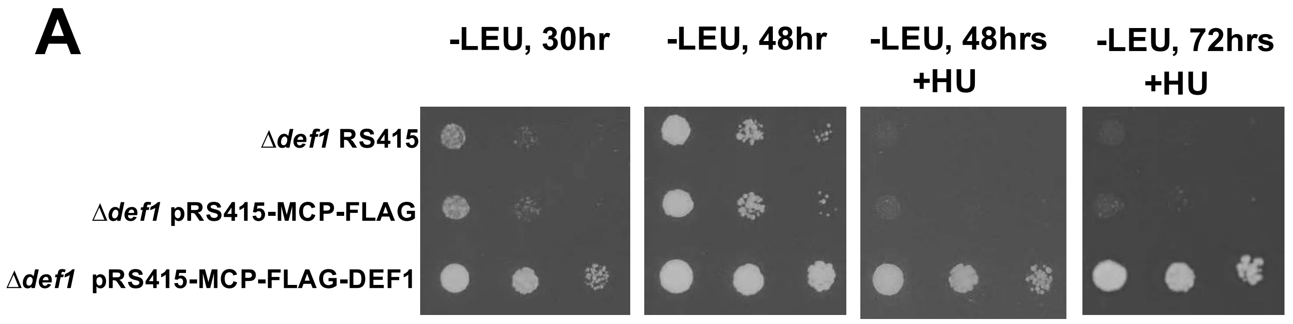


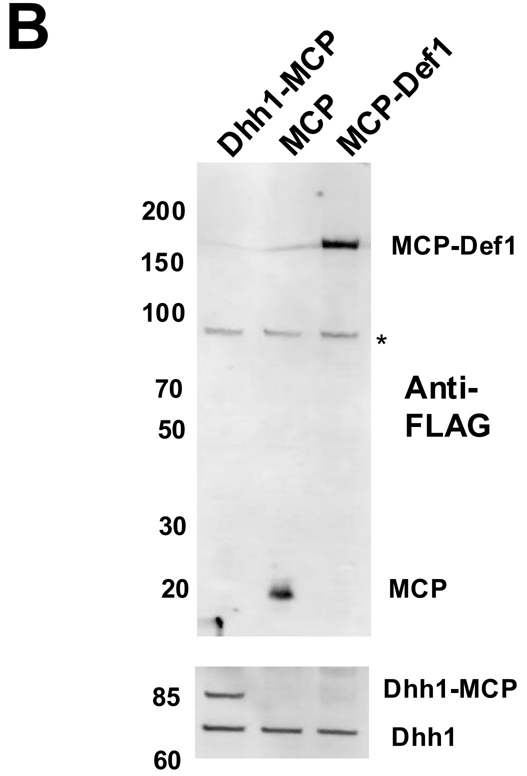


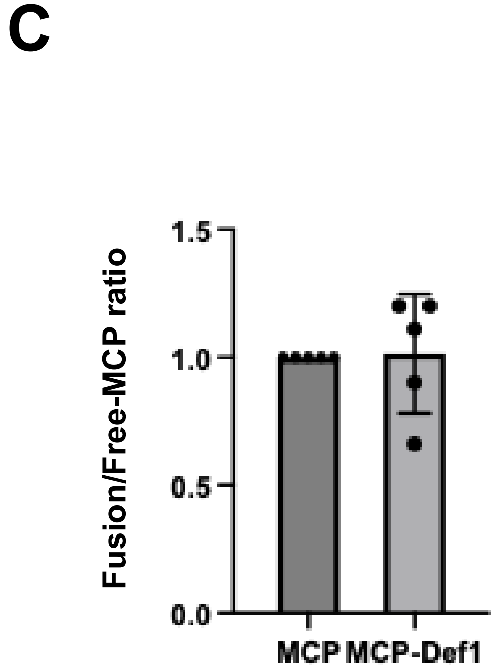

Supplementary figure 6

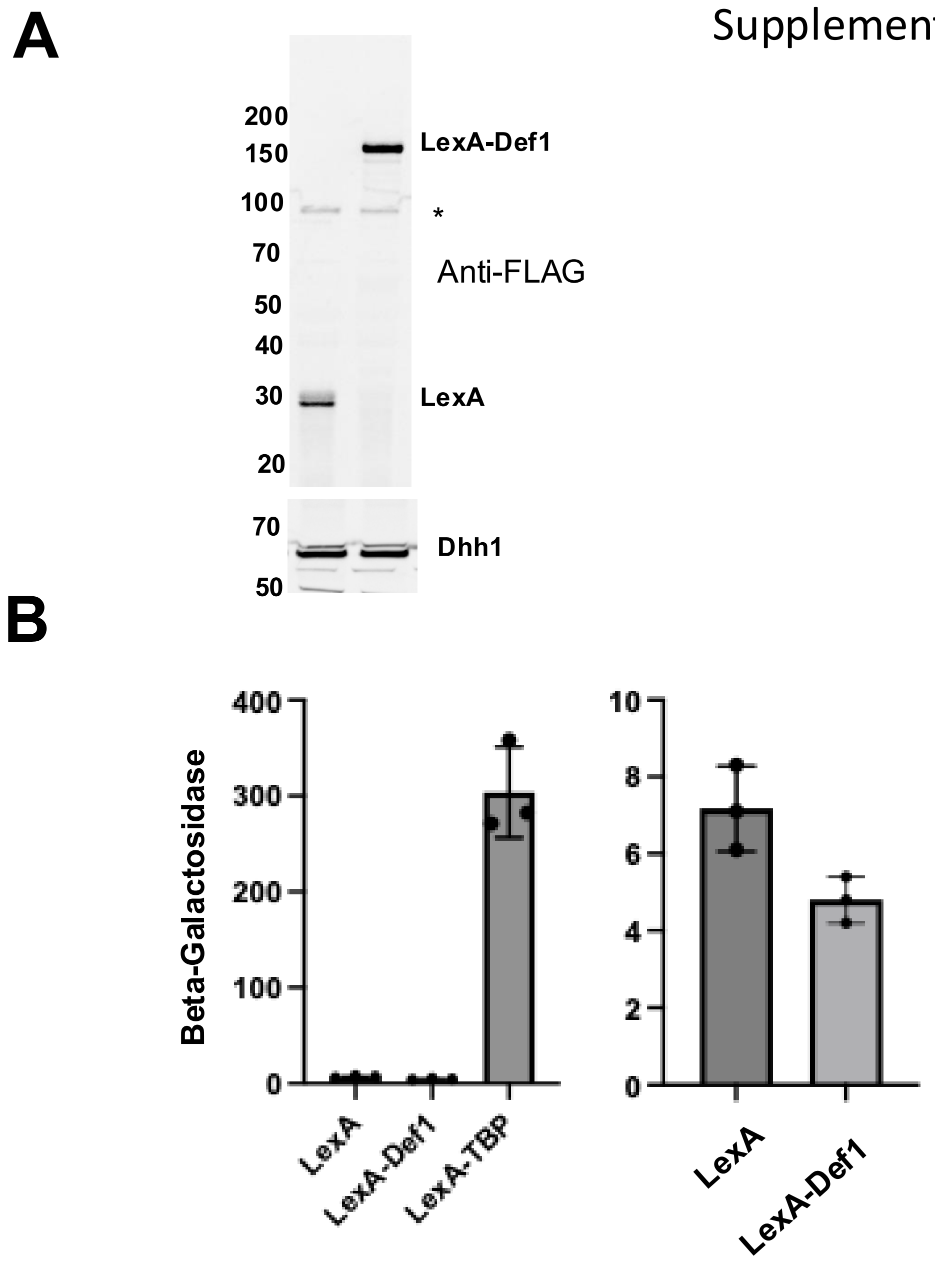

**Supplemental figure legends**

**Supplemental Figure 1.** (A). Correlation of biological replicates of total RNA-seq reads. The reads (FPKM) from biological replicates of each strain were plotted in a scatterplot. Pearson correlation coefficients were determined for the reads. ShinyGO 0.8 was used to identify GO terms for KEGG pathways of transcripts with an FC>2, and $p_{adj}<0.01$

**Supplemental Figure 2. Correlation of RATE-seq reads between biological replicates.** RATE-seq reads from biological replicates at each labeling time point (6 min, 9 min, 12 min, 24 min, 90 min) were plotted on a scatterplot. Pearson correlation coefficients between replicates were calculated and displayed on the respective plots.

**Supplemental Figure 3. Comparison of the median RNA half-life from this and other studies.** The differences in the methods and conditions are shown for comparison, since this can affect the value estimated from the studies. (MILLER et al. 2011; MUNCHEL et al. 2011; NEYMOTIN et al. 2014; PRESNYAK et al. 2015; BAUDRIMONT et al. 2017; CHAN et al. 2018; ALALAM et al. 2022).

**Supplemental figure 4. Turbo-ID shows that Def1 labels proteins in the cell.** (A). Spot test of wild type (BY4742) and Def1-TID-3HA cells (JR2200). Serial dilutions of saturated cultures were spotted onto YAPD and YAPD+ 75 mM hydroxyurea (HU). (B). Western blot of free TID-3HA expressed from the *CHA1* promoter (JR1951) and Def1-TID-3HA (JR2200). Dhh1 is used as a loading control. The asterisk indicates a protein cross-reacting with the HA antibody. (C). A representative streptavidin-pull-down. Proteins were eluted with SDS-PAGE loading buffer containing 3 mM biotin, separated on SDS-PAGE gels, and transferred to nitrocellulose. Biotinylated proteins were then detected using streptavidin-HRP.

**Supplemental Figure 5: Supporting information for mRNA and promoter tethering experiments.** (A). Complementation assay. A *def1*∆ strain (JR2204) was transformed with the plasmids indicated on the left. Cells were serially diluted and spotted on the media indicated above the panel. HU was used at a concentration of 75 mM. (B). Representative western blot of cells expressing MCP, MCP-Def1 and Dhh1-MCP. The upper blot was probed with anti-FLAG (M2). The lower blot was detected using anti-Dhh1 antibody. Asterisk marks a cross-reacting band. A 3X-FLAG epitope was incorporated into the MCP and MCP-Def1 fusion proteins. Dhh1-MCP is not FLAG tagged and is not detected by the M2 antibody. (C). Quantification of the expression of the MCP fusion proteins. The proteins were detected using anti-FLAG antibodies. Total amounts of protein were controlled for by blotting for Taf14. The average expression of the MCP-Def1 fusion protein, relative to the amount of free MCP, which was set to 1.0. Average with standard deviations (N=5) is presented.

**Supplemental Figure 6. Promoter tethering assay.** (A). Anti-FLAG western blot. Dhh1 is the loading control (B). LexA- transcription activation reporter assay. (left) A Beta-galactosidase reporter gene driven by the *GAL1* core promoter containing 8 lexA binding sites (GOLEMIS et al. 2001) was co-transformed with plasmids expressing “free” LexA , LexA-Def1, and LexA-TATA-binding protein (TBP). LexA-TBP is a positive control, which drives expression by increasing PIC formation (CHATTERJEE AND STRUHL 1995) Three biological replicates were analyzed. (right) Plotting only the free LexA and LexA-Def1 derivatives to display the details on Beta-galactosidase expression.
